# Effects of Antiretroviral/ Cotrimoxazole Treatment on Metabolic Markers are Linked to Immune Reconstitution and Depletion of the Gut Protist *Blastocystis*

**DOI:** 10.64898/2026.11.24.769077

**Authors:** John O’Connor, Angela Sofia Burkhart Colorado, Nichole M Nusbacher, Janine Higgins, Charles Preston Neff, Suzanne Fiorillo, Thomas B Campbell, Margaret Borok, F. Kathryn Boyd, John Sterrett, Brent E Palmer, Catherine Lozupone

## Abstract

**Background:** HIV remains a global health threat, with many people with HIV (PWH) residing in sub-Saharan Africa (SSA). Despite effective antiretroviral therapy (ART), metabolic complications are common. In this longitudinal study in Zimbabwe, we enrolled ART-naïve, ART-treated, and healthy control (HC) participants and investigated factors associated with metabolic health.

**Approach:** All cohorts were sampled 24 weeks apart, with the ART-naïve cohort sampled before and after treatment with ART and cotrimoxazole prophylaxis. Metabolic markers, including high and low-density lipoprotein cholesterol (HDL-C and LDL-C), triglycerides, and glucose, were related to blood inflammatory cytokines and immune cell populations and the fecal microbiome composition and its functional potential, while controlling for confounding factors including diet. Mediation analyses were conducted for causal inference.

**Results:** HDL-C was significantly lower in ART-naïve compared to ART-treated PWH and HCs and increased longitudinally with treatment. Increases in HDL-C correlated with immune reconstitution with reductions in inflammation identified as a significant mediator. However, treatment was also associated with increased triglyceride and glucose concentrations. Depletion of intermediate monocytes mediated triglyceride increases. Loss of the intestinal protist *Blastocystis*, a known target of cotrimoxazole, correlated with loss of bacterial diversity and higher fasting glucose concentration. Microbial functional potential, notably utilization of the phosphotransferase system, correlated with improved metabolic health across cohorts.

**Conclusions:** Treatment with ART and cotrimoxazole and concomitant immune reconstitution was associated with reversal of compromised HDL-C levels in PWH. However, the potential effects of treatment on commensal microbes, such as *Blastocystis,* may contribute to treatment-induced metabolic effects in PWH.

**Importance:** The gut microbiome’s influence on metabolic health in PWH in SSA is not well understood. We provide insights into this relationship while considering HIV- and ART-associated immune dysfunction/reconstitution and other potential confounders. Standard of care prophylactic cotrimoxazole treatment protects against opportunistic infections in PWH in SSA and largely spares anaerobic intestinal commensals. However, cotrimoxazole directly targets the intestinal protist *Blastocystis,* which is common in parts of SSA and has been associated with higher gut microbiome diversity and improved metabolic health. Our results suggest that the decreased inflammation occurring with treatment of PWH reverses HIV-associated reductions in HDL-C, a healthy form of cholesterol. *Blastocystis* associated with both higher gut microbiome diversity and lower fasting glucose, and was significantly reduced with treatment, suggesting that *Blastocystis* loss may be a driver of poor metabolic health markers that arise with cotrimoxazole prophylaxis.

## Introduction

HIV type 1 (HIV-1) infects approximately 39 million individuals globally, with many residing in sub-Saharan Africa (SSA).^1^ With HIV infection, CD4+ T depletion and immune dysfunction occur^2^ along with increased prevalence of metabolic syndrome (MetS), characterized by insulin resistance, hypertension, and dyslipidemia. This contributes to increased risk of cardiovascular disease and early mortality.^3,4^ While lifespan of people with HIV (PWH) has been improved by effective antiretroviral therapy (ART), this has been associated with increased prevalence of dyslipidemia, diabetes, and altered fat distribution.^5^ Understanding the pathophysiology of metabolic co-morbidity in PWH can inform treatments.

The underlying causes of HIV-associated metabolic dysfunction are multifactorial and can include immune activation/inflammation,^6^ direct effects of HIV proteins R (Vpr) and Nef,^7–12^ and drug-specific ART toxicities.^13^ Older classes of ART drugs, including protease inhibitors and reverse transcriptase inhibitors, can have particularly strong effects.^14–16^ First-line ART regimens often used in SSA have been associated with weight and metabolic changes.^17^ Efavirenz (EFV), which was used as a first line therapy until its replacement in 2019 by dolutegravir (DTG), has been associated with increases in fasting plasma total cholesterol, high-density lipoprotein cholesterol (HDL-C), low-density lipoprotein cholesterol (LDL-C), triglycerides, and serum glucose levels.^18^ Furthermore, the change to DTG has been associated with increased weight gain and increased risk of hypertension.^19^

In addition to ART, prophylactic cotrimoxazole is often standard of care in SSA since it reduces the risk of opportunistic infections in PWH. Both ART and cotrimoxazole can induce microbiome disturbances,^20–26^ but it is unknown whether these disturbances are related to metabolic abnormalities in PWH. One way that gut microbiome differences have the potential to affect the metabolic health in PWH is through disruptions to intestinal barrier integrity and as drivers of chronic peripheral immune activation and inflammation.^27–32^ Poor metabolic health in PWH has previously been associated with increased barrier translocation markers and with gut microbiome signatures such as increased Desulfovibrionaceae and decreased *Butyrivibrio* and *Coprococcus*.^27,33,34^ Furthermore, *in vitro* stimulations of human immune cells with microbial communities isolated from feces have supported that fecal microbiomes that induce high immune-activation are associated with high peripheral immune activation in PWH.^35^ Understanding relationships between microbiome composition, immune activation, and metabolic health markers in SSA has been under-explored. Prior studies investigating relationships between the microbiome and metabolic health have focused on gut bacterial composition, with less attention given to microbial eukaryotes or microbiome functional potential.^36^ Understanding eukaryotic populations may be particularly important in SSA, as a recent large-scale study linked the intestinal protist *Blastocystis*, a common gut commensal in the developing world, with favorable metabolic health.^37^

In this longitudinal study of PWH and controls conducted in rural and urban centers in Zimbabwe, we assessed relationships between markers of metabolic health, including fasting concentrations of circulating glucose, high-density lipoprotein cholesterol (HDL-C), low-density lipoprotein cholesterol (LDL-C), and triglycerides, and several host and microbial factors, including peripheral immune activation and inflammation, the prokaryotic (16S) and eukaryotic (18S) components of the gut microbiome, and gut microbiome functional potential. We accounted for potential confounders including diet and age. Individuals were assessed at baseline and 24 weeks after initiating ART with efavirenz/lamivudine/tenofovir disoproxil fumarate (EFV/3TC/TDF) plus cotrimoxazole (Naïve). Two additional groups were also recruited and assessed at both timepoints; 1) PWH already receiving EFV/3TC/TDF and cotrimoxazole (Exp) and 2) healthy control (HC) participants. Data from this cohort has been used previously to demonstrate differential treatment-associated immune recovery in urban and rural settings and the influence of dietary elements on immune and microbial differences by HIV and treatment status.^22,38^ We hypothesized that treatment-associated changes in metabolic markers would be influenced by the degree of immune recovery and changes to the gut microbiome, finding potential effects of immune reconstitution and loss of intestinal *Blastocystis* on metabolic marker changes that occurred with treatment.

## Results

### Cohort and Baseline Confounders

Baseline comparisons of measures across cohorts are displayed in **Table 1**. No differences in sex were observed, but the Exp group was older and had lower Body Mass Index (BMI). As expected, Exp had lower viral load and higher CD4+ T cell counts.

**Table 1:** Study Population: Baseline data, split by cohort, presented as n, median (interquartile range) or n (%), unless otherwise stated. Statistics determined by Kruskal-Wallis with Dunn’s post-hoc test for continuous variables and chi-square test for categorical variables, Naïve=ART-naïve, Exp=ART-experienced, HC=healthy controls, BMI=body mass index, IQR=inter-quartile range, LDL-C=low-density lipoprotein cholesterol, HDL-C=high-density lipoprotein cholesterol, IL-6=Interleukin-6, CRP=C reactive protein, *=approximate estimates assuming individuals started in June of the year they indicated starting ART.

| **Variable** | **Naïve** | **Exp** | **HC** | **Exp vs HC** | **Exp vs Naive** | **HC vs Naive** |
|---|---|---|---|---|---|---|
| Sample Number | 65 | 33 | 40 |  |  |  |
| **Arm** |  |  |  | **1.000** | **1.000** | **1.000** |
| Urban (Count (%)) | 32 (49.2%) | 17 (51.5%) | 20 (50%) | NA | NA | NA |
| Rural (Count (%)) | 33 (50.8%) | 16 (48.5%) | 20 (50%) | NA | NA | NA |
| **Sex** |  |  |  | **1.00** | **1.000** | **1.000** |
| Female (Count (%)) | 37 (56.9%) | 17 (51.5%) | 21 (52.5%) | NA | NA | NA |
| Male (Count (%)) | 28 (43.1%) | 16 (48.5%) | 19 (47.5%) | NA | NA | NA |
| **Demographics** |  |  |  |  |  |  |
| Age (years) (Median (IQR)) | 34 (26-39) | 46 (42-51) | 35.5 (28-41.25) | **<0.001** | **<0.001** | 0.203 |
| **Clinical Characteristics** |  |  |  |  |  |  |
| BMI (Median (IQR)) | 22.5 (20-24) | 20.2 (18.7-23) | 22.45 (20.6-24.92) | **0.020** | 0.093 | 1.000 |
| CD4 Count (Median (IQR)) | 257 (123-358) | 412 (249-575) | NA (NA-NA) | **NA** | **0.001** | NA |
| Plasma HIV-1 RNA (copies/mL) (Median (IQR)) | 26491 (3985-71968) | 19 (19-19) | NA (NA-NA) | NA | **0.004** | NA |
| Approximate Duration on ART in Years (Median (IQR)) * | 0.79 (0.60-0.87) | 7.41(6.35-10.05) | NA (NA-NA) | NA | **<0.001** | NA |
| Duration on Cotrimoxazole in Months (Median (IQR)) | 0.03(0-0.23) | 784 (60-108) | NA (NA-NA) | NA | **<0.001** | NA |
| **Metabolic Markers** |  |  |  |  |  |  |
| HDL-C (mg/dL) (Median (IQR)) | 41.5 (30-49) | 58 (55-69) | 50 (41.5-62) | 0.203 | **<0.001** | **<0.001** |
| LDL-C (mg/dL) (Median (IQR)) | 82.5 (60.75-101) | 84 (67-112) | 82.5 (68.5-116.5) | 1.000 | 1.000 | 1.000 |
| Triglycerides (mg/dL) (Median (IQR)) | 75.5 (57.75-96.25) | 78 (58-120) | 71.5 (52.25-88.25) | 0.318 | 1.000 | 1.000 |
| Glucose (mg/dL) (Median (IQR)) | 83 (78-87.25) | 87 (81-91) | 81 (75.75-88.5) | 0.078 | 0.189 | 1.000 |
| **Inflammatory Markers** |  |  |  |  |  |  |
| IL-6 (pg/mL) (Median (IQR)) | 2.74 (1.09-5.8) | 1.47 (1.03-3.3) | 1.45 (0.8-2.07) | 1.000 | 0.175 | **0.015** |
| CRP (mg/L) (Median (IQR)) | 2.67 (0.61-5.88) | 1.95 (0.47-4.68) | 2.06 (0.77-3.73) | 1.000 | 1.000 | 1.000 |
| **16S Microbiome Alpha Diversity (N)** |  |  |  |  |  |  |
| Shannon Diversity (Median (IQR)) | 6.03 (5.68-6.35) | 5.51 (4.98-6.24) | 6.33 (5.77-6.64) | **0.004** | 0.054 | 0.601 |
| Faith Diversity (Median (IQR)) | 70.9 (54.81-83.69) | 58.69 (46.27-74.29) | 78.87 (57.81-96.02) | **0.035** | 0.480 | 0.418 |
| **18S Microbes Detected (N)** |  |  |  |  |  |  |
| Blastocystis (N (%)) | 27 (41) | 8 (24.2) | 24 (60) | **0.014** | 0.428 | 0.305 |

### ART and Cotrimoxazole Increased HDL-C in the Naïve Group, but also Increased Glucose and Triglycerides

We explored relationships between metabolic markers (fasting levels of blood glucose, HDL-C, LDL-C, and triglycerides) and HIV and treatment status. At baseline, the Naïve cohort had lower HDL-C, which is generally considered to be a healthy form of cholesterol ^39–41^, compared to the Exp and HC groups (**Table 1**). Baseline fasting blood glucose concentrations were trending higher in Exp compared to HCs suggesting a slight treatment effect that was not statistically significant. However, most remained in the healthy range of fasting glucose (under 100 mg/dL) as determined using previously established thresholds^42^. There were no baseline differences across cohorts in LDL-C or triglycerides (**Table 1**; **Figure 1B**). To select confounders, we used multiple variable selection algorithms to identify relationships between metabolic markers and age, BMI, gender, location (urban/rural), and dietary intake, including total dairy, snacks, drinks, grains, protein, fast food, fruit, and vegetables (**Supplementary Figure 1A and 1B**). Variables selected from the best performing models (lowest R-squared) were included in downstream linear models (**Figure 1A**); These included dietary components (fast food, grain, and protein), BMI, which positively correlated with LDL-C, age, which positively correlated with LDL-C and triglycerides, and location, in which individuals in the urban location had increased glucose and triglyceride concentrations. These analyses were consistent with the overall trends, with HIV status associated with decreased HDL-C and treatment associated with HDL-C improvement.

**Figure 1:**
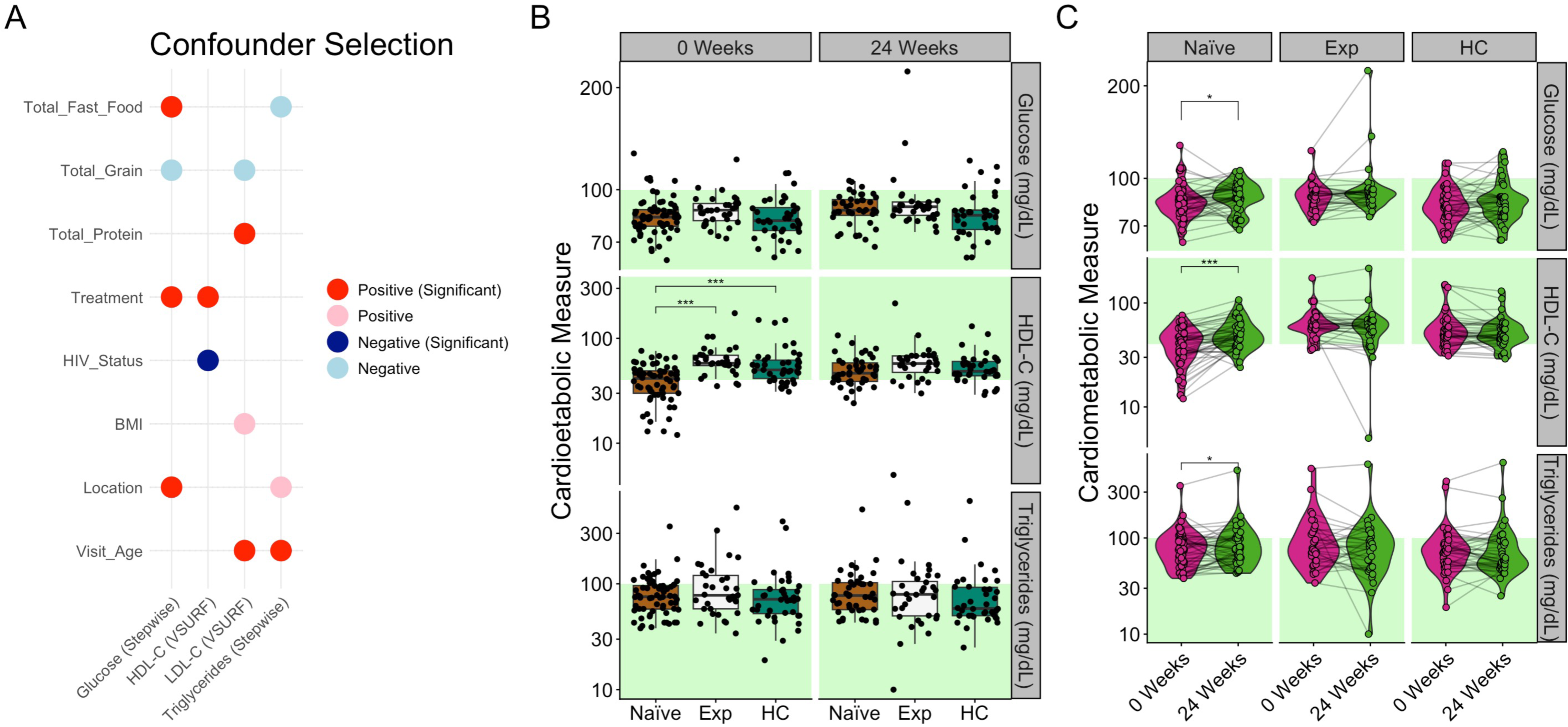
Metabolic markers Related to Age, HIV, and Treatment: (A) Coefficients of linear regressions comparing metabolic markers (x-axis) from baseline demographic and dietary predictors (y-axis) that were selected by different variable selection approaches, which include lasso VSURF, and stepwise. Significance was determined by t-test for linear regression with p-values then FDR adjusted across all predictors. Significant features in the final model are in bold colors and had FDR adjusted p-values (q) under 0.05. Each column represents one linear model for the listed marker, with variable selection approach in parentheses. Circles represent features included in the model after variable selection. For categorical variables, red is associated with positive HIV status (HIV status), individuals receiving treatment (Treatment), and individuals in the rural cohort (Location) where blue indicates the opposite. (B) Box plots of plasma metabolic markers split by cohort and timepoint. Statistical significance across three groups was determined by Kruskal-Wallis and pairwise comparisons made using Dunn’s post-hoc test with Bonferroni correction for multiple comparisons. (C) Violin plots of plasma metabolic markers across the study period with spaghetti plots joining values from the same subject overlaid; brackets indicating significance as determined by paired Wilcoxon; *-p≤0.05, **-p≤0.01, ***-p≤0.001, ****-p≤0.0001. Healthy ranges for each measure by previously established thresholds^43^ indicated with green shading.

We also evaluated changes in metabolic markers that occurred in the Naïve cohort with 24 weeks of treatment. Consistent with cross-sectional analysis, HDL-C and glucose had statistically significant increases, and triglycerides additionally increased (**Figure 1C**). This included a median increase of 11.5 mg/dL HDL-C, 8.0 mg/dL triglycerides, and 6.0 mg/dL glucose in the Naïve group, suggesting both beneficial (HDL-C) and detrimental (triglycerides and glucose) effects. Because ART and cotrimoxazole were co-administered per standard of care, observed treatment effects reflect their combined impact and cannot be attributed to one component alone.

### Reduced Inflammation Associated with Metabolic Marker Changes

We next sought to understand relationships between inflammation and metabolic health markers, using measurement of circulating C reactive protein (CRP) and IL-6 concentrations. As we have previously reported,^22^ IL-6 was higher in the Naïve cohort compared to HC participants and to a lesser extent Exp participants at baseline, but CRP did not differ (**Table 1**). IL-6 concentration decreased with ART/cotrimoxazole treatment, but CRP exhibited no statistically significant change (**Supplementary Figure 2A**).^22^ To assess the relationship between inflammatory and metabolic markers, linear mixed effects models (LMEMs) were performed accounting for confounding variables identified with variable selection (**Figure 1A**). We performed these analyses while including all participants, or just the individuals in the Naïve cohort to focus on longitudinal treatment effects. CRP concentration positively correlated with triglyceride concentration and IL-6 concentration negatively correlated with HDL-C concentration when considering all participants, consistent with a detrimental effect of inflammation on metabolic markers. However, IL-6 concentration was negatively related to LDL-C and triglyceride concentrations (**Supplementary Figure 2B**), which is the opposite of the expected direction. When only considering the Naïve cohort, IL-6 concentration was also negatively related to HDL-C concentration, and mediation analysis revealed that decreased IL-6 with 24 weeks of treatment was a significant mediator of an increase in HDL-C concentration (p=0.016) (**Supplementary Figure 2C,D**).

### HDL-C Increase with ART was associated with Immune Reconstitution

Immune cell populations determined by flow cytometry differed cross-sectionally across the 3 cohorts as previously described.^22^ After 24 weeks of treatment, the Naïve participants exhibited increased CD4+ T cell percentage and decreased CD8+ T cell percentage, decreased T cell activation (CD4+ and CD8+ T cells expressing HLA-DR and CD38), decreased T cells expressing PD-1) and decreased CD14+ intermediate monocytes (**Figure 2A**). In analyses conducted across all study participants and within the Naïve group only, LMEMs did not identify any relationships between specific immune cell populations and any metabolic markers after correcting for multiple tests. However, mediation analysis revealed that the treatment-associated increase in triglycerides was mediated by decreased CD14+ intermediate monocytes (p=0.004) (**Figure 2B**). Immune reconstitution of the Naïve cohort to an immune profile similar to that of the HC and Exp participants with 24 weeks of treatment is shown in the ordination plot of the flow cytometry data (**Figure 2C and D**). HDL-C was positively correlated with PC2 (**Figure 2E and F**), which was itself positively associated with markers of immune reconstitution, including increased CD4+ T cell percentages and decreased T cell activation and exhaustion markers.

**Figure 2:**
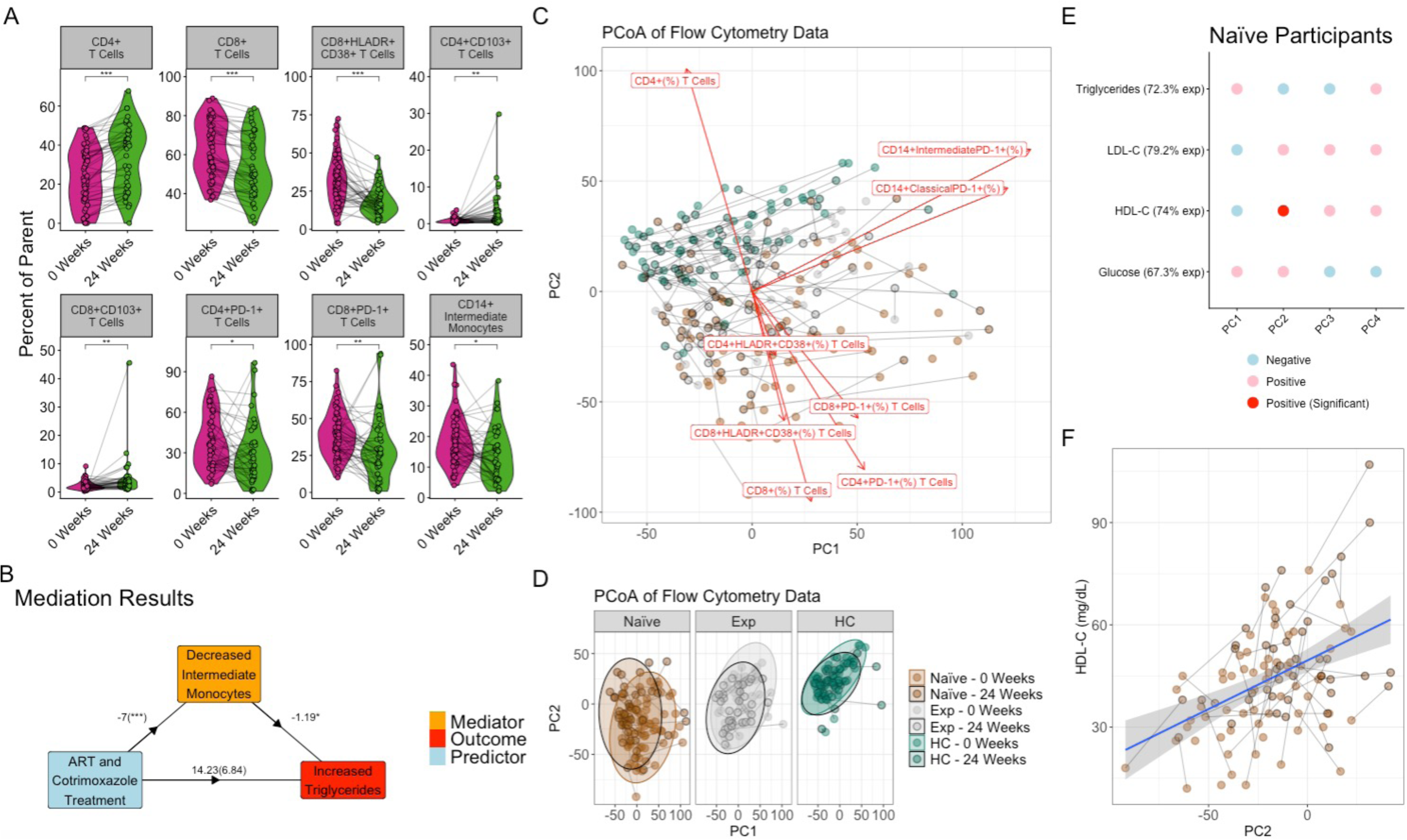
(A) Violin plots overlaid with spaghetti plots of immune cell populations that significantly changed upon ART treatment with significant determined by Friedman test; (B) Mediation results diagram, boxes indicate predictors and markers, statistics on arrows indicate the linear model coefficient comparing the input (beginning of the arrow) to the output (end of the arrow) with asterisks indicating significance by model t-test, parenthesis indicates the coefficient with the mediator (orange box) incorporated into the model (C) Ordination biplot of immune cell populations colored by cohort and timepoint; (D) Ordination plot of immune cell populations split by cohort with 95% confidence ellipses; black outlines indicate visit 2 data (E) Coefficients of LMEMs comparing metabolic markers (y-axis) to ordination axes with confounders (x-axis) in ART naïve participants with significance determined by t-test for linear regression with p-values then FDR adjusted, significance determined by adjusted p-values under 0.05 (F) Scatter plot of PC2 values (x-axis) HDL-C values (y-axis) in naïve cohort, with lines connecting the same subject and black outlines indicating visit 2 data; ; *-p≤0.05, **-p≤0.01, ***-p≤0.001, ****-p≤0.0001

### Metabolic Markers Related to Microbiome Composition

We next sought to understand the relationship between treatment-associated changes to the fecal bacterial microbiome and metabolic markers using previously generated 16S rRNA gene sequencing data. As reported previously,^22^ treatment was associated with decreased α-diversity, with the Exp group having lower Shannon diversity and Faith phylogenetic diversity (PD) compared to the HC group at baseline, and the Naïve cohort having a significant drop in both α-diversity measures with treatment (**Supplementary Figure 3A,B**). *Lachnoclostridium* decreased in Naïve participants following treatment.^22^

In all study participants, LMEMs with selected confounders revealed that Faith PD was negatively associated with HDL-C (**Supplementary Figure 3C and 3D**) but no significant relationships between specific genera and metabolic markers were found after correcting for multiple comparisons. To explore relationships with overall microbiome composition, we calculated weighted UniFrac distances^44^ between pairs of samples and performed PCoA (**Supplementary Figure 4B-E)**. LMEMs comparing Weighted UniFrac PC axes to metabolic markers, revealed that PCo3, which was related to microbiomes with higher levels of *Succinivibrio* and *Prevotella*, positively correlated with HDL-C concentration (**Supplementary Figure 4F and G**).

In the Naïve cohort, changes in metabolic markers had no significant relationship with changes in α-diversity using LMEM or in mediation analysis. Because *Lachnoclostridium* reduced with treatment and has been associated with poor metabolic health,^45^ mediation analysis was performed but yielded no significant results. However, in the Naïve cohort, we found that HDL-C concentration positively correlated with *Anaerobutyricum hallii* group (R^2^=0.77, Coefficient=0.051, padj=0.025) and negatively correlated with *Candidatis soleaferrea* (R^2^=0.78, Coefficient=-0.078, padj=0.025) and that fasting glucose concentration negatively correlated with *Rothia* (R^2^=0.85, Coefficient=-0.021, padj=0.027).

### The Impact of treatment on Gut *Blastocystis*

To evaluate relationships between eukaryotic members of the intestinal microbiome and metabolic markers, we generated new 18S rRNA sequencing data. The most frequently detected eukaryotic microorganism at baseline was *Blastocystis*, being detected in 44.3% of samples in the population overall. At baseline, this *Blastocystis* prevalence differed by cohort (**Figure 3A**), and location (**Figure 3B**). *Blastocystis* was both more often detected (p-adj=0.014) and had a higher relative abundance (p-adj=2.20×10^-4^) in the HC cohort compared to the Exp cohort (**Figure 3A,C**) and more often detected (p-value=1.53×10^-4^) and had a higher relative abundance (p-value=0.001) in the rural compared to the urban samples (**Figure 3B,D**). Furthermore, in the Naïve group, 24 weeks of treatment reduced *Blastocystis* relative abundance (p-value=0.047), and this was driven by a reduction in the rural area (**Figure 3E**). The Naïve group had decreased prevalence and relative abundance of *Blastocystis* compared to the HC cohort at week 24 but not at baseline, again reflecting the drop with treatment (**Figure 3A**). Interestingly, at both baseline and 24 weeks, *Blastocystis*-high (over the median abundance) had lower fasting glucose compared to the *Blastocystis*-low group (below the median abundance) (**Figure 3F**) and *Blastocystis* abundance negatively correlated with fasting glucose at baseline (Spearman: Rho=-0.209, p-value=0.014) and 24 weeks (Spearman: Rho=-0.213, p-value=0.023) (**Figure 3G**). *Blastocystis*-high samples also had significantly higher α-diversity at both timepoints (**Figure 3H**), and significantly different microbiome composition as determined by weighted UniFrac (PERMANOVA:R^2^=0.042,p=0.001) (**Figure 3I**).

**Figure 3:**
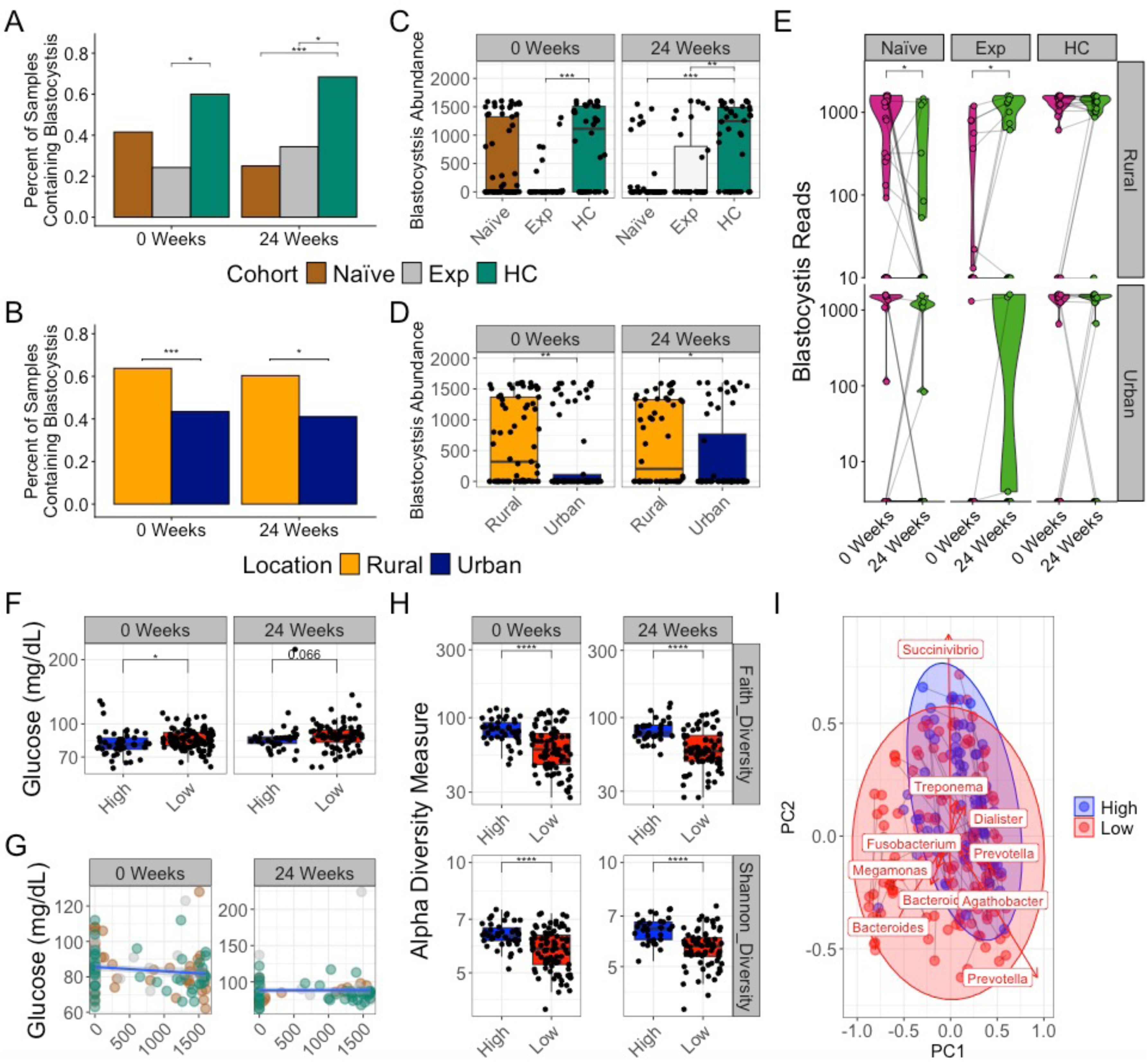
(A,B) Bar plot of the prevalence of *Blastocystis* in fecal samples at baseline and visit 2, (A) split by cohort and (B) split by location, with brackets indicating significant differences across groups with Chi-squared test (C,D) Boxplot of abundance of *Blastocystis* rarified reads across (C) cohort and (D) location, split by study visit, with brackets indicating significant differences across groups using (C) Kruskal-Wallace test and Bonferroni correction and (D) Wilcoxon signed-rank test; (E) Longitudinal violin plots with overlaid spaghetti plots connecting participants across the study period split by cohort and location, with brackets indicating significance as determined by Friedman test; (F) boxplot of plasma glucose in *Blastocystis*-high and *Blastocystis*-low samples split by study visit, with brackets indicating significant differences across groups using Wilcoxon signed-rank test (G) scatter plot of plasma glucose versus abundance of *Blastocystis* reads with trend line established by linear models with 95% confidence range, statistics represent Statistics obtained by Spearman correlations; (H) Boxplots of alpha diversity metrics across *Blastocystis*-high and *Blastocystis*-low samples, with brackets indicating significant differences across groups using Wilcoxon signed-rank test; (I) Ordination biplots of 16S results using the weighted UniFrac distance metric colored by *Blastocystis* category; *-p≤0.05, **-p≤0.01, ***-p≤0.001

### Comparison to Functional Potential as Determined by Shotgun Metagenomic Sequencing

Shotgun metagenomic sequencing was performed on 32 baseline samples from the rural participants, including 16 from Naïve, 8 from Exp, and 8 from HC, to perform a previously reported analysis of relationships between gut microbiome composition and diet using samples selected to represent individuals consuming a spectrum of diets.^38^ Here we used these data to further explore relationships between microbiome functional potential and metabolic markers. While no significant relationships between specific functional genes and metabolic markers were detected by linear models after FDR correction, Gene Set Enrichment Analysis (GSEA) revealed several gene sets enriched and depleted with metabolic markers when accounting for confounders (**Figure 4A-D**). Notably, the phosphotransferase system (PTS), which includes a set of genes responsible for detection and translocation of complex carbohydrates and downstream fermentation, was positively related to HDL-C concentration, but negatively associated with triglyceride and LDL-C concentrations.

**Figure 4:**
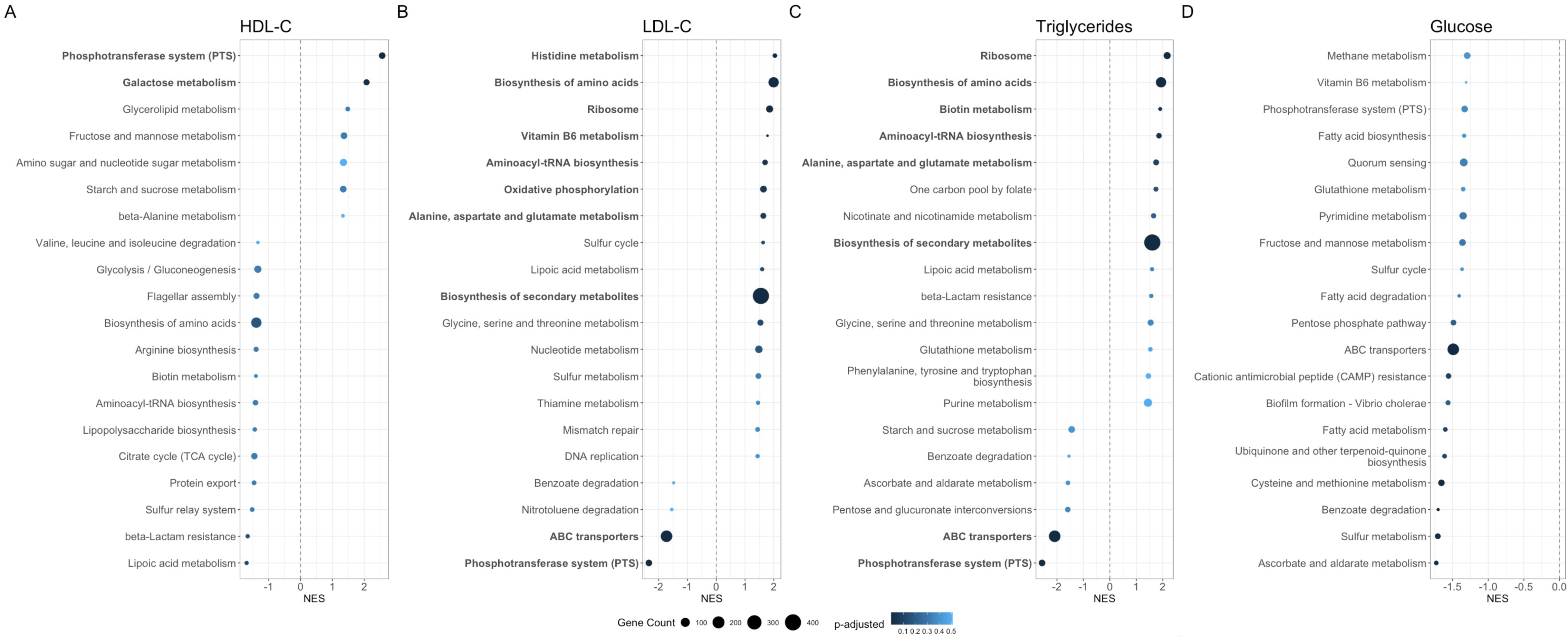
Dot plots of gene set enrichment analysis results of the 20 most enriched or depleted gene sets (y-axis) related to (A) HDL-C, (B) LDL-C, (C) Triglycerides, and (D) Glucose, sorted by net enrichment score (NES) (x-axis), size of dots indicate the gene set size, color indicates the adjusted p-value, and bolding indicates significantly enriched or depleted gene sets (adjusted p value <0.05). Enrichment analysis was performed on by ranking coefficients obtained from LMEMs, the formula being *Metabolic Marker∼Gene Abundance+Confounders+(1|PID)*, where PID is the participant ID and the confounders include those established by the baselin variable selection.

## Discussion

We found that untreated PWH in Zimbabwe had lower HDL-C concentration than HC or treated PWH and that treatment with ART and cotrimoxazole resulted in HDL-C improvement, but fasting glucose and triglyceride concentrations also small increases, suggesting both potential beneficial and detrimental effects on metabolic health. ART was given together with cotrimoxazole, which is commonly used in SSA to prevent infections in PWH.^46,47^ It is thus difficult to disentangle ART versus cotrimoxazole effects. While HDL-C improvement was related to immune reconstitution and IL-6 reduction, increased fasting glucose concentration was related to reduced intestinal *Blastocystis* which a target of cotrimoxazole, and increased triglyceride concentration was related to reduced CD14+ intermediate monocytes. These findings indicate that drivers of metabolic health in PWH on treatment are complex and multifactorial, as has been reviewed previously^48^.

### HDL-C with HIV and Treatment

HDL-C is generally considered to be beneficial for cardiometabolic health,^39,40^ as it is responsible for returning cholesterol to the liver from peripheral tissues, thereby preventing atherosclerosis and heart disease.^41^ In fact a median improvement of 11.5 HDL-C is associated with approximately 30% reduced risk of coronary heart disease as established by the Framingham Heart Study^49^. Our results are consistent with previous studies that also showed reduced HDL-C concentration with untreated HIV infection,^50,51^ increased HDL-C concentration with diverse ART regimes,^52,53^ and a relationship between HDL-C increases and CD4+ T cell recovery with ART.^54^ The overserved mediating effect of reduced inflammation (IL-6) on HDL-C is consistent with studies demonstrating an inverse relationship between IL-6 and HDL-C in other disease contexts.^55,56^ However, one prior study has also associated cotrimoxazole use with lower incidence of low-HDL in PWH on ART, indicating that direct effects of cotrimoxazole could also be a contributing factor.^57^

Treatment-associated microbiome changes observed here, including loss in α-diversity and *Lachnoclostridium*, were not correlated to HDL-C changes. Since prior studies have shown gut microbiome α-diversity to positively correlate with HDL-C concentration,^58,59^ the loss of microbiome diversity with treatment would be expected to have the opposite effect on HDL-C than we observed. However, we did find significant relationships between HDL-C concentration and microbiome composition and function unrelated to the taxa that changed with treatment. For instance, *Anaerobutyricum (formally Eubacterium) hallii*^60^, correlated positively with HDL-C concentration in the Naïve cohort. *Anaerobutyricum (formally Eubacterium) hallii* has been shown to have beneficial effects on insulin resistance via butyrate production in a murine diabetes model,^61^ and also to have higher expansion in MetS individuals receiving microbiome infusions from lean donors who had better cardiometabolic improvement.^62^ We also found HDL-C concentration to positively correlate with the PTS pathway and galactose metabolism in GSEA analysis of shotgun metagenomic data of baseline samples. PTS is a complex system of microbial genes involved in carbohydrate uptake transport and phosphorylation,^63^ suggesting beneficial effects of high-affinity carbohydrate scavenging of microbiomes. This functional profile likely supports the production of downstream fermentation products, such as butyrate, which has been associated with epithelial barrier strength^64^ and immune homeostasis.^65^ Butyrate does induce ATP-binding cassette subfamily A member 1 (ABCA1) ^66^which contributes to the biosynthesis of HDL-C^67^, suggesting fermentation could be contributing to circulating HDL-C concentrations. However, it is worth noting that the relationship between microbiome functional potential and metabolic health is context dependent, as Turnbaugh et al. found that microbiomes with elevated carbohydrate utilization specifically galactose metabolism are associated with obesity and confer weight gain in gnotobiotic mice^68^, suggesting microbiome diet utilization can have both positive and detrimental effects on metabolic health depending on dietary intake.

### Potential Mediation of Detrimental Treatment Effects on Metabolic Markers by ART Directly

We observed modest increases in fasting glucose and triglyceride concentrations with treatment. These markers were not higher in Naïve individuals compared to HCs at baseline, suggesting that these are negative effects on metabolic health related specifically to treatment. ART has been previously associated with MetS-related^13^ and diabetic clinical manifestations in PWH^69–71^, including in SSA specifically.^72,73^ Overall, NNRTI-based therapies such as the one used here have not had as strong effects as INSTIs (dolutegravir and bictegravir), which have been associated with weight gain, and early Protease Inhibitors (PIs), which have been associated with lipodystrophy/fat redistribution.^69,74,75^ However, plasma levels of EFV, which is a component of the EFV/3TC/TDF combination therapy used in this cohort, have been associated with increased lipids and serum glucose concentration,^18^ which could have partially contributed to the increases that we observed. EFV has largely been replaced by DTG in first line therapies since 2019 because of this and other side-effects.^17^

### Potential Mediation of Increased in Fasting Glucose Concentration with Treatment by Blastocystis Loss

ART was given together with cotrimoxazole, which is commonly used in SSA to prevent infections in PWH.^46,47^ Cotrimoxazole has been shown to alter gut microbiome in mice and humans,^25,76^ but is known to have limited direct effects on anaerobic intestinal commensals;^77^ Cotrimoxazole’s mechanism of action is to inhibit folate biosynthesis, and most gut commensals can acquire dietary folate. Indeed, one controlled study of cotrimoxazole intake actually showed increased α-diversity.^25^ However, *Blastocystis* is highly sensitive to cotrimoxazole,^78^ and this intestinal protist has been strongly linked to gut microbiome α-diversity in SSA^76,77,78,79^ and elsewhere.^80,81^ Our data support that the loss of diversity with treatment may be influenced by cotrimoxazole-driven loss of *Blastocystis.* This protist was detected at high levels, particularly in rural study participants, consistent with other studies conducted in SSA.^36,82–84^ Interestingly, we had previously reported that α-diversity decreases with treatment only occurred in the rural and not urban participants of this study,^85^ and similarly here, we show that a decrease in *Blastocystis* also only occurred in the rural area where prevalence was high. This may suggest that the loss of α-diversity with treatment could be driven by *Blastocystis* loss rather than direct action of cotrimoxazole. This may explain inconsistent results of cotrimoxazole on α-diversity across studies. For instance in our prior work in Cameroon, where *Blastocystis* prevalence is low,^37^ α-diversity loss was linked to type of ART and not observed with certain ART types even though individuals were taking cotrimoxazole.^86^

Interestingly, a recent large-scale analysis of *Blastocystis* and metabolic markers showed a strong association with *Blastocystis* presence and improved metabolic markers.^37^ This is consistent with our finding that *Blastocystis* correlated with reduced plasma glucose. In a large-scale shotgun metagenomic study of participants enrolled in the Personalized Responses to Dietary Composition Trial (PREDICT 1),^87^ *Blastocystis* presence was associated with lower post-prandial glucose concentrations. Our results however do suggest that *Blastocystis* could serve as a target for future mechanistic investigations, especially in certain areas of SSA where it is more prevalent.^37^ One unanswered question is whether *Blastocystis* may influence metabolic health by promoting bacterial diversity or specific compositions. However, except for a negative relationship between fasting glucose and *Rothia* in the Naïve cohort, microbiome associations for glucose were not strong. It is also worth noting that while *Blastocystis* could help maintain plasma glucose homeostasis, it is also possible lower glucose could be causing *Blastocystis* outgrowth, as clinical trials in type 2 diabetes have shown that improved glucose homeostasis results in elevated *Blastocystis*^88^, so additional studies are required to investigate causality.

### Intermediate monocytes and triglycerides

Changes in microbiome composition were not associated with the higher triglyceride concentrations with treatment, suggesting microbiome independent mechanisms. Baseline triglyceride concentrations were related, however, to microbiome functional pathways, including a negative relationship with the PTS pathway discussed above in relation to HDL-C concentrations. While treatment conferred immune reconstitution in our Naïve group, it also reduced CD14+ intermediate monocytes which mediated triglyceride increases. This is consistent with past studies showing that CD14+ intermediate monocytes are lower in HIV-positive women receiving the same EFV/3TC/TDF combination compared to HIV seronegative controls,^89^ and that reduced CD14+ intermediate monocytes are associated with reduced viral load^90^. This suggests that, on ART, this reduced monocyte subtype is linked to the lowering of viral burden. CD14+ intermediate monocytes are also significant producers of IL-6^91^ indicating their reduction could be contributing to reduced IL-6. However, with regard to triglycerides, intermediate monocytes have been associated with uptake of triglyceride rich lipoproteins.^92^ Therefore, their reduction with treatment could be reducing triglyceride clearance, leaving higher levels in plasma. Despite this effect, the overall impact of reduced CD14+ intermediate monocytes on cardiometabolic health is probably complex as other studies have shown a positive relationship between intermediate monocytes and cardiovascular complications,^93–95^ and HIV infection has also been shown to alter atherosclerosis-associated intermediate monocyte transmigration.^96^ However, intermediate monocytes are very heterogeneous and can exhibit different functions in different contexts. ^91,97,98^

### Effects of Other Factors

Although the main interest of our study was the effect of HIV infection and treatment on metabolic health markers, we identified several other influencing factors that we treated as confounders in our statistical models. Our finding that age associated positively with triglyceride and LDL-C concentrations is consistent with studies showing increased prevalence of dyslipidemia and MetS with age.^4,99–101^ Age remains a significant predictor of metabolic health within this Zimbabwe cohort despite varied HIV infection and treatment status. While HIV can exacerbate the inflammatory and metabolic effects of aging, strikingly age was more predictive of elevated LDL-C and triglyceride concentrations than either HIV infection or treatment status, despite their known contribution to poor metabolic health.^4,13,102–104^

Diet had a less pronounced but significant effect on metabolic health markers. The positive relationship between frequency of fast food intake and glucose concentration observed across all participants is also consistent with prior studies.^105,106^ The positive relationship between LDL-C concentration and protein intake was surprising, but could be due to increased poultry and beef intake, which were included in that category^38^, and have been associated with increased LDL-C concentrations.^107^

Our study is not without limitations. Our Exp and Naïve groups were receiving cotrimoxazole and ART simultaneously, limiting our ability to delineate between the two. Also, while mediation analysis does provide mechanistic insight into *potential* causality, further controlled studies are required for confirmation.

## Materials and Methods

### Study Design and Recruitment

This study was approved by multiple ethics and regulatory committees, including the Joint Research Ethics Committee for University of Zimbabwe Faculty of Medicine and Health Sciences (UZFMHS), Parirenyatwa Hospital (JREC), the Research Council of Zimbabwe, Medical Research Council of Zimbabwe (MRCZ), and the Colorado Multiple Institutional Review Board (COMIRB; 16-1441). Written informed consent was obtained before any research related activities. Both English and Shona versions of the informed consent form were available. Participants were recruited from Zimbabwe at two locations, the urban Mabvuku Polyclinic in Harare and the rural Mutoko District Hospital. Only adults over 18 years of age were recruited. The Naïve cohort were PWH who had less than 10 days of ART prior to study entry with EFV/3TC/TDF. Study participants also had prophylactic cotrimoxazole as part of standard of care treatment. The Exp cohort were PWH who were treated with EFV/3TC/TDF for a minimum of 12 months prior to entry. This cohort also had cotrimoxazole with standard of care treatment. Exclusion criteria were use of antibiotics (except cotrimoxazole) within 2 months of enrollment, pregnancy, or a BMI greater than 29 kg/m^2^.

### Collection of Participant Information and Samples

Study participants completed questionnaires, in which information regarding age, HIV status, BMI, and gender was collected. Trained interviewers provided a Food Frequency Questionnaire (FFQ) in which dietary information was collected as has been reported previously^38^. Stool samples were collected within 24 hours after visit 1 and visit 2 in a fecal collection kit and aliquoted into an OmniGene Gut Collection System for DNA preservation. Venipuncture was used to obtain fasting blood samples, which were collected in BD vacutainers. Blood was centrifuged to separate plasma, while white blood cells (WBC) from the remaining blood were isolated immediately by density centrifugation for flow cytometry. Plasma and fecal samples were initially stored at -80°C at the Infectious Diseases Research Laboratory in the Internal Medicine Department at the University of Zimbabwe Faculty of Medicine and Health Sciences in Harare. They were then shipped on dry ice to the University of Colorado Anschutz Medical Campus and stored at -80°C upon arrival.

### Measurement of Plasma Markers

Enzyme-linked immunosorbent assays (ELISAs) were used to measure blood inflammatory markers, including IL-6 and CRP as previously described^22,38^. Enzymatic assays to measure HDL-C, LDL-C, triglycerides, and glucose were conducted at the Clinical and Translational Research Center (CTRC) at the University of Colorado Anschutz Medical campus ^22,38^.

### Flow Cytometry

Flow cytometry was used to characterize blood immune cell as previously described but was expanded to include additional populations.^22,38^ Red blood cells (RBCs) were lysed using 1x RBC lysis buffer. Cells were then washed twice with staining buffer containing phosphate buffered saline (PBS), 2% Bovine Serum Albumin (BSA), and 1 mM Ethylenediaminetetraacetic Acid (EDTA). Cells were surface stained with BV785-labelled anti-CD3 antibody, PerCP/Cy5.5-labelled anti-CD4 antibody, BV421-labelled anti-CD8 antibody, BV605-labelled anti-CD38 antibody, Pe-labelled anti-CD103 antibody, Pe-Cy7-labelled anti-HLA-DR antibody, BV510-labelled anti-CD14 antibody, APC-H7-labelled anti-CD16 antibody, and FITC-labelled anti-PD-1 antibody or appropriate fluorescence minus one (FMO) controls. Cells were then washed again twice with staining buffer followed by fixation in 1% formaldehyde.

Cells were analyzed on a BD LSR Fortessa Flow Cytometer. Data were analyzed using FlowJo. The cell populations included in downstream analysis, were CD14+ classical monocytes, which included CD14+ HLA-DR+ classical monocytes, CD14+ PD-1+ classical monocytes, CD14+ intermediate monocytes, which included CD14+ HLA-DR+ intermediate monocytes, CD14+ PD-1+ intermediate monocytes, and CD3+ T cells, which included CD4+ T cells, CD4+ CD103+ T cells, CD4+ PD-1+ T cells, CD4+ HLA-DR+ CD38+ T cells, CD8+ T cells, CD8+ CD103+ T cells, CD8+ PD-1+ T cells, and CD8+ HLA-DR+ CD38+ T cells.

### 16S and 18S Sequencing

Fecal DNA was extracted, amplified, and pooled as previously described^22,38^. Sequencing data was obtained, demultiplexed, denoised, and filtered as described^22,38^. For 18S data, samples were rarified to 1,603 reads and presence of *Blastocystis* was determined by the detection of any reads taxonomically classified to the *Blastocystis* genus. For the 16S data, samples were rarified to 16,645 reads. The QIIME 2 SATé-Enabled Phylogenetic placement (SEPP) plugin was used to make a phylogenetic tree for diversity measures^108^. Shannon diversity, Faith’s Phylogenetic Diversity^109^, and weighted UniFrac distances^44^ were calculated with QIIME 2.

### Shotgun Metagenomic Sequencing

The same extracted DNA was also used for shotgun sequencing on a subset of baseline samples. Sequencing was performed at the Genomics core at the University of Colorado. Qubit and TapeStation 4200 were used to determine gDNA purity, quantity, and size distribution. Covaris was used to mechanically sheer an input of 55 ng of gDNA targeting 300-400bp of DNA products. DNA-seq libraries were generated using the Ovation Ultraflow System V2 kit. 150bp paired-end sequencing reads were generated using a NovaSeq X plus sequencer with a 600 million paired-end reads per sample target depth. Demultiplexing was performed using bcl2fastq. Trimmomatic was used for trimming raw sequencing reads, and FastQC and MultiQC were used for quality assessment.^110,111^ Host reads were filtered using hostile with the masked human-t2t-hla-argos985 reference genome.^112^ Reads were functionally characterized using the Human Microbiome Project (HMP) Unified Metabolic Analysis Network (HUMAnN) available through bioBakery 3.^113^

### Quantification and Statistical Analysis

To compare continuous variables across groups we used Kruskal-Wallis tests with Dunn’s post hoc tests for more than two groups and Wilcoxon signed-rank test for two groups. Categorical comparisons across groups were assessed using chi-square tests. To assess the relationships between predictive variables on metabolic outcomes we used linear models, with metabolic outcomes log transformed. Confounders were identified using variable selection by random forest (VSURF) using the VSURF R Package^114^, Lasso using the brms R package^115^, or stepwise approaches with the stats R package^116^. For cross sectional data at set timepoints, linear modelling was performed using the stats R package. For longitudinal data, linear mixed effects models were performed treating participant ID as a random variable using the lme4 R package^117^. To assess the significance of specific variables in the linear models we used a regression t-test using the summary function in R. Gene set enrichment was performed on genes ranked by fold change (categorical comparisons) or coefficients of models (continuous comparisons) using the clusterProfiler^118^ R package only including gene sets ranging from 10 to 500 genes.

## Data Availability Statement

Analysis code for this project is available on GitHub (https://github.com/jboconnor13/Zimbabwe-Cardiometabolic-Analysis). The 16S rRNA, 18S rRNA, and shotgun sequence data are both available on ENA (project PRJEB66206, accession ERP151280, project PRJEB105653, accession ERP186804, and project PRJEB95999, accession ERP178750, respectively).

## Acknowledgments

We would like to thank our study participants for their time and dedication to the study. We would also like to thank Mercia Mutimuri and Patricia Gambiza for support in recruitment and sample collection from study participants, Francis Jaji for study coordination support, the staff of the Infectious Diseases Research Laboratory, University of Zimbabwe for sample processing and storage, and the staff of the Mutoko District Hospital and Mabvuku Polyclinic.

